# Pericarp Pigmentation Correlates with Hormones and Intensifies with Continuation of Bud Sport Generations from ‘Red Delicious’

**DOI:** 10.1101/336263

**Authors:** Wen-Fang Li, Juan Mao, Shi-Jin Yang, Zhi-Gang Guo, Zong-Huan Ma, Mohammed Mujitaba Dawuda, Cun-Wu Zuo, Ming-Yu Chu, Bai-Hong Chen

## Abstract

Bud sport mutants of apple (*Malus domestica* Borkh.) trees with a highly blushed colouring pattern are mainly caused by the accumulation of anthocyanins in the pericarp. Hormones are important factors modulating anthocyanin accumulation. However, a good understanding of the interplay between hormones and anthocyanin synthesis in apples, especially in mutants at the molecular level, remains elusive. Here, physiological and comparative transcriptome approaches were used to reveal the molecular basis of pericarp pigmentation in ‘Red Delicious’ and its mutants, including ‘Starking Red’, ‘Starkrimson’, ‘Campbell Redchief’ and ‘Vallee spur’, which were designated G0 to G4, respectively. Pericarp pigmentation gradually proliferated from G0 to G4. The anthocyanin content was higher in the mutants than in ‘Red Delicious’. The activation of early phenylpropanoid biosynthesis genes, including *ASP3*, *PAL*, *4CL*, *PER*, *CHS*, *CYP98A* and *F3’H*, was responsible for anthocyanin accumulation in mutants. In addition, IAA and ABA had a positive regulatory effect on the synthesis of anthocyanins, while GA had the reverse effect. The down-regulation of *AACT1*, *HMGS*, *HMGR*, *MVK*, *MVD2*, *IDI1* and *FPPS2* involved in terpenoid biosynthesis influences anthocyanin accumulation by positively regulating transcripts of *AUX1* and *SAUR* that contribute to the synthesis of IAA, *GID2* to GA, *PP2C* and *SnRK2* to ABA. Furthermore, MYB and bHLH members, which are highly correlated (*r*=0.882–0.980) with anthocyanin content, modulated anthocyanin accumulation by regulating the transcription of structural genes, including *CHS* and *F3’H*, involved in the flavonoid biosynthesis pathway.

## INTRODUCTION

Bud sport is a somatic mutation occurring in the shoot cells of perennial fruit trees and an important source of discovery of new cultivars or strains that are superior to the parent (Petit and Hampe 2006; El-Sharkawy *et al*. 2015). Although the genetic background of these mutants is nearly identical to that of their parents (Nwafor *et al*. 2014; Otto *et al*. 2014), epigenetic changes have been recognized, such as causing fruit colour alteration in apple (*Malus domestica* Brokh.) (Xu *et al*. 2012; El-Sharkawy *et al*. 2015). Pericarp colour is a key appearance and nutrition quality attribute of apple fruit (Willams and Benkeblia 2018). Anthocyanins are secondary metabolites that contribute to the colours of fruits (Meng *et al*. 2016). Pigmentation in the skin of apple fruit varies among cultivars and is influenced by environmental factors, including temperature (Arrizabalaga *et al*. 2018) and the level of sunlight irradiation (Cominelli *et al*. 2007; Honda and Moriya 2018). Furthermore, hormones are likely to be important factors that modulate light-dependent anthocyanin accumulation (Jeong *et al*. 2004; Carvalho *et al*. 2010; Loreti *et al*. 2010). In summary, exploring the molecular mechanisms of hormones and anthocyanin synthesis in apple fruit and its bud sport mutants is crucial to research on pigment accumulation and plant somatic mutation.

Previous studies have shown that auxin (IAA), cytokinin (CTK), gibberellins (GA), jasmonate acid (JA), abscisic acid (ABA) and ethylene (ETH) interact in controlling anthocyanin biosynthesis (El-Kereamy *et al*. 2003; Jeong *et al*. 2004; Loreti *et al*. 2010; Liu *et al*. 2014; Ji *et al*. 2014). In addition, the identification and functional characterization of MYB and bHLH transcriptional factors revealed that they play a role in autonomously mediated structural gene transcription. These factors include chalcone isomerase (*CHI*), chalcone synthase (*CHS*), flavonol synthase (*FLS*), leucoanthocyanidin reductase (*LAR*), flavonoid 3’-hydroxylase (*F3’H*) and anthocyanidin reductase (*ANR*), which are involved in the anthocyanin biosynthesis pathway (Deluc *et al*. 2008; Gonzalez *et al*. 2008; Telias *et al*. 2011; Petroni and Tonelli 2011; An *et al*. 2012). MYB proteins are characterized by two imperfect repeats of the DNA-binding motifs R2 and R3 (Ramsay and Glover 2005), and bHLH proteins are characterized by the basic helix-loop-helix domain, which is responsible for sequence-specific DNA binding (Massari and Murre 2000).

The publication of the apple reference genome (Daccord *et al*. 2017) and the development of new tools for transcriptomics have facilitated recent advances in the genome-wide analysis of dynamic gene expression during pericarp development (El-Sharkawy *et al*. 2015; Massonnet *et al*. 2017). The strategies of hormone and anthocyanin synthesis are often applied without a full understanding of the effect at the molecular level, with the exception of a few studies that have correlated biochemical and physiological outcomes with transcriptomic changes (Jeong *et al*. 2004; Carvalho *et al*. 2010; Loreti *et al*. 2010). Apple cultivar ‘Red Delicious’ (*M. domestica*) is the most frequently captured sport apple variety that is usually selected on the phenotypic basis of spur type and intense red fruit colour. The cultivar’s four continuous generation mutants, namely, ‘Starking Red’, ‘Starking Red’, ‘Starkrimson’, ‘Campbell Redchief’ and ‘Vallee Spur’, have been screened. Therefore, these five strains were selected and analysed using a comparative transcriptome combined with physiological and biochemical characteristics to expound the relationships between hormone and anthocyanin synthesis on apple pericarp pigment accumulation.

## MATERIALS AND METHODS

### Plant material

‘Red Delicious’ is the most frequently captured sport apple variety, featuring four continuous generation mutants. ‘Starking Red’ is a bud sport from ‘Red Delicious’ and a typical representative of the first generation. The second generation is ‘Starkrimson’, which is a bud sport from the first-generation ‘Starking Red’. The fourth generation, ‘Vallee Spur’, is a bud sport of the third generation, ‘Campbell Redchief’, which is bud sport of ‘Starkrimson’. Mature apple fruit of these five cultivars range from having red vertical stripes to being completely red (see Figure 1A).

**Figure 1.**
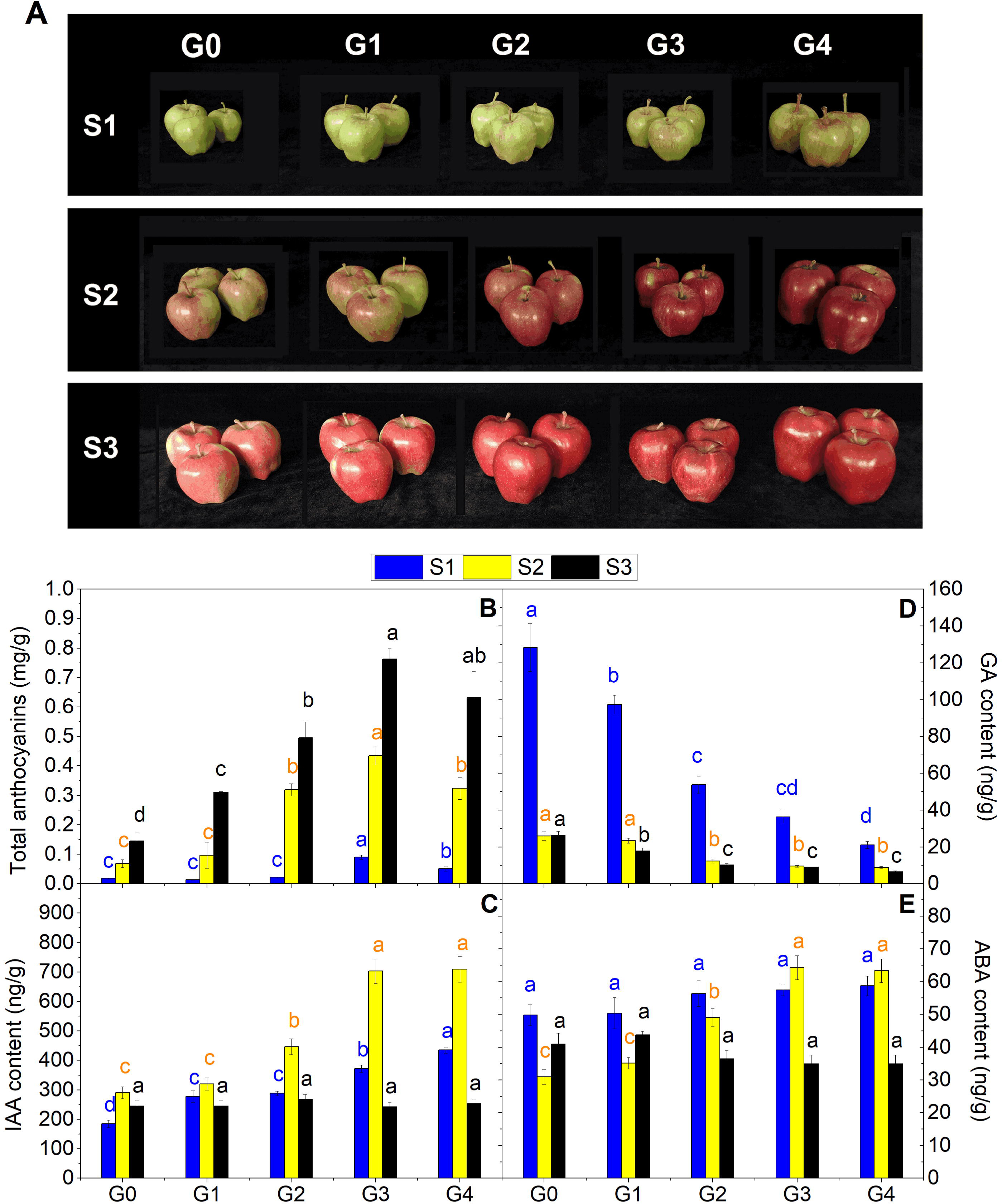
A, Close-up views of ‘Red Delicious’ and its four generation mutants (‘Starking Red’, ‘Starkrimson’, ‘Campbell Redchief’ and ‘Vallee spur’), named G0 to G4, at three developmental stages (S1–S3) used for anthocyanin quantification, transcriptome profiling and qRT-PCR. B, The changes in total anthocyanin concentrations in the pericarp of the five strains at S1 to S3. Changes in endogenous hormone levels, including IAA (C), GA (D) and ABA (E), in the pericarp of the five strains at S1 to S3. Values are means ± SE. Different lower case letters indicate significant differences among the five strains (*P*=0.05).

Fruit pericarp samples of ‘Red Delicious’ and its four continuous generation mutants (‘Starking Red’, ‘Starkrimson’, ‘Campbell Redchief’ and ‘Vallee spur’) were named G0 to G4 and collected in 2017 from 12-year-old trees grown in apple demonstration gardens at Tianshui, China. Briefly, 20−30 fruits from each of the five strains were sampled at three developmental stages, i.e., 5 August (S1), 25 August (S2), and 14 September (S3) (see Figure 1A). Stages S1, S2 and S3 are equivalent to the pre-veraison, veraison and fruit maturity stages for commercial harvest, respectively. In different experiments, pericarp samples from 6 fruits per replicate with three independent biological replicates were collected. All samples were collected at the same time of day (9–10 _AM_), immediately frozen in liquid nitrogen and stored at −80 °C for further analysis of anthocyanin contents, endogenous hormone contents and gene expression profiles (qRT-PCR). In addition, samples from S2 were used for RNA-seq analysis.

### Anthocyanin quantification

Lyophilized apple pericarp samples were finely ground, and approximately 500 mg of powdered samples was homogenized in 10 mL of methanol with 1% HCl. The homogenate was transferred to a calibration test tube with a constant volume of 20 mL and kept for 20 min at 4 °C with shaking under dark conditions. Then, the samples were filtered through a 0.2 µm polyethersulfone (PES) filter (Krackeler Scientific, Inc., Albany, NY, USA) and analysed using a TU-1900 double beam UV-visible spectrophotometer (Beijing Purkinje General Instrument Co. LTD). Anthocyanin levels were calculated by dividing the absorbance by the coefficient of regression (0.0214) acquired by standard scale measurements.

### Hormone content measurement

A total of 1.0 g of each lyophilized apple pericarp sample was ground quickly after liquid nitrogen was added and combined with 10 mL of 80% chromatographic pure methanol. Each sample was washed three times with solvent, transferred into a test tube, and stored in a refrigerator at 4 °C overnight in the dark. Then, the samples were centrifuged for 20 min under refrigerated conditions at 4 °C. Supernatant fluid was transferred into a new centrifuge tube. The extract was concentrated, and the methanol was volatilized at 40 °C by rotary evaporation to obtain 2 mL of concentrate. The evaporation bottle wall was then washed continuously with 50% methanol, and the volume was raised to 10 mL with 50% chromatographic pure methanol. The fluid for testing was filtered through a 0.22 μm organic membrane.

The determination method was performed with different concentrations of IAA, GA, and ABA, standard samples, which were used to construct a standard curve. The standard samples were purchased from Sigma Company, and the external standard curve and quantitative methods were performed for the measurements. The apparatus used for high-performance liquid chromatography (HPLC) was the LC-20AD system (Shimadzu, Kyoto, Japan) equipped with a Zorbax Eclipse Plus C18 column (4.6 mm × 250 mm × 5.0 μm, Agilent, Palo Alto, CA, USA) and an SPD-20A UV detector. The mobile phase was chromatographic methanol and 0.6% iced acetic acid. The flow velocity was 1.0 mL/min, the wavelength was 254 nm, and the column temperature was 25 °C.

### RNA extraction

Total RNA was extracted from approximately 200 mg of lyophilized apple pericarp samples ground in liquid nitrogen using the RNase-Free DNase Set (Qiagen, Valencia, CA, USA) and then cleaned with the RNeasy Mini Kit (Qiagen). RNA quality and quantity were determined using a Pultton P200 Micro Volume Spectrophotometer (Pultton Technology Limited).

### Library preparation and sequencing

The 5 triplicate samples (5 varieties at S2) yielded 15 nondirectional cDNA libraries with a total of 68.18 million reads (Table 1), which were prepared from 3.0 µg of total RNA using the NEBNext, Ultra^TM^ RNA Library Prep Kit (NEB, USA). RNA quality and quantity were assessed using a NanoDrop spectrophotometer and an Agilent 2100 spectrophotometer (Agilent Technologies, CA, USA). Library fragments were purified with an AMPure XP system (Beckman Coulter, Beverly, USA), and the quality was assessed on an Agilent Bioanalyzer 2100 system. Index-coded samples were clustered on a cBot cluster generation system using the TruSeq PE Cluster Kit v3-cBot-HS (Illumina Inc., San Dego). After cluster generation, library preparations were sequenced on an Illumina HiSeq 2000 platform, in which 125 bp paired-end reads were generated.

### Analysis of sequencing results: mapping and differential expression

Raw reads were cleaned by removing adapter sequences, reads containing ploy-N, and low-quality sequences (Q < 20). Clean reads were aligned onto the apple reference genome (https://iris.angers.inra.fr/gddh13/) (Daccord *et al*. 2017). New transcripts were identified from TopHat alignment results using the Cufflinks v2.1.1 reference-based transcript assembly method. An average of 83.95% of reads were mapped for each sample (Table 1). For annotations, all novel genes were searched against the NCBI non-redundant protein sequence database (Nr), Swiss-Prot, Gene Ontology (GO) database, Cluster of Orthologous Groups of proteins (COG), protein family (Pfam), and Kyoto Encyclopedia of Genes and Genomes (KEGG) database using BLASTx with 10^-5^ as the E-value cut-off point; sequences with the highest similarities were retrieved. After amino acid sequences of new genes were predicted, they were searched against the Pfam database using HMMER software, and annotation information of these new genes was obtained.

The DESeq (2010) R package was utilized to detect differentially expressed genes (DEGs). The false discovery rate (FDR) was used to identify the *P-*value threshold in multiple tests (Benjamini and Hochberg, 1995). An FDR < 0.01 and fold changes ≥ 2 were used as screening criteria; an absolute value of log_2_ (fold change) with reads per kb per million reads (FPKM) ≥ 1.0 was used as a threshold to determine significant DEGs (Mortazavi *et al*. 2008).

### Functional analysis of differentially expressed genes (DEGs)

Functional enrichment analysis, including GO and KEGG, was performed to identify DEGs that were significantly enriched in GO terms or metabolic pathways. GO enrichment analysis of DEGs was implemented by the GOseq R package, in which gene length bias was corrected. GO coupled with *KS*<0.01 was considered significantly enriched by DEGs (see Table S1). KOBAS software was used to test the statistical enrichment of different expression genes in KEGG pathways. Pathways with a *Q*-value ≤ 0.05 were defined as genes that displayed significant levels of differential expression (see Table S2).

### Common expression pattern clustering analysis of DEGs

The different expression patterns of DEGs among the five strains were analysed using the R language, Cluster package, Biobase package, and *Q*-value package. The DEGs with a common expression trend were divided into a data set, which was expressed as a model map. The distance measure used was Euclidean distance, and the clustering method was K-means clustering or hierarchical clustering.

### Correlation analysis

A correlation matrix was prepared using SPSS statistical software and Pearson’s correlation coefficient as the statistical metric. The analysis was performed using the anthocyanin content at S2 and the FPKM average of each candidate DEG. Correlation values were converted to distance coefficients to define the height scale of the dendrogram.

### Quantitative real-time PCR validation of RNA-Seq data

Quantitative reverse transcription RT-PCR analysis DNase-treated RNA (2 µg) was reverse transcribed in a reaction volume of 20 µl using PrimerScript^TM^ RT reagent Kit with gDNA Eraser (Takara, Dalian, China). Gene-specific primers were designed using Primer Express software (Applied Biosystems) (see Table S3). Quantitative reverse transcription PCR (qRT-PCR) assays were performed using 20 ng of cDNA and 300 nM of each primer in a 10 µl reaction with SYBR Green PCR Master Mix (Takara, Dalian, China). Three biological and three technical replicates for each reaction were analysed on a LightCycler^®^ 96 SW 1.1 instrument (Roche). The amplification program consisted of one cycle of 95 °C for 30 s, 40 cycles of 95 °C for 5 s, and melting analysis at 60 °C for 34 s, followed by one cycle of 95 °C for 15 s, 60 °C for 60 s, and 95 °C for 15 s. Transcript abundance was quantified using standard curves for both target and reference genes, which were generated from serial dilutions of PCR products from corresponding cDNAs. Transcript abundance was normalized to the reference gene *MdGADPH*, which showed high stability across the different apple genotypes and tissues used in this study. Relative gene expression was normalized by comparing with G0 expression and analysed using the comparative 2^-ΔΔCT^ method (Livak and Schmittgen 2001).

### Data analysis

Data regarding the anthocyanin content and the relative expression level of specific genes were analysed by ANOVA, and treatment means were separated by Duncan’s multiple range test at *P* < 0.05 with the aid of SPSS statistical software. For correlation analysis, the Pearson correlation coefficient (*r*) was calculated, and a two-tailed test was carried out.

### Data availability

Supplemental materials available at FigShare. Dataset 1 contains the mean of the normalized expression value per transcript (FPKM fragments per kilobase of mapped reads) of the 5-sample at S2. Dataset 2 contains DEGs identified as commonly up-regulated or down-regulated in each pairwise comparison of ‘Red Delicious’ and its four generation mutants. Dataset 3 contains gene composition of the six clusters identified using gene expression clustering analysis. Dataset 4 contains gene composition and mean FPKM value of the phenylpropanoid/flavonoid biosynthesis pathway. Dataset 5 contains correlation analysis of anthocyanin content at S2 and MYB transcriptional factors. Dataset 6 contains correlation analysis of anthocyanin content at S2 and bHLH transcriptional factors. Dataset 7 contains gene composition and mean FPKM value of the terpenoid biosynthesis pathway. Dataset 8 contains gene composition and mean FPKM value of the plant hormone signal transduction.

## RESULTS

### Pericarp pigmentation increased with bud sport generation and maturation

Visual inspection of apple pericarp colour during development revealed that the G2, G3 and G4 strains began colouring at S1, with the most visibly intense red colouring occurring in G3 and G4 (see Figure 1A). Pigmentation of these five strains progressively advanced to much higher levels from S1 to the subsequent stage S3, resulting in red fruit at maturity. In addition, red coloration gradually proliferated from G0 to G4 during the three stages.

Consistent with visual inspection, the analysis of apple pericarp anthocyanin contents showed that the levels of total anthocyanins in the five strains were initially low and sharply increased with maturation (from S1 to S3) (see Figure 1B). Relative to that at G0, the total anthocyanin level in fruit skin was ~0.78-, ~1.20-, ~5.02- and ~2.85-fold higher in G1, G2, G3 and G4 at stage S1, respectively. At S2, the level in fruit skin was ~1.41-, ~4.65-, ~6.33- and ~4.72-fold higher in G1, G2, G3 and G4, respectively. Finally, at S3, the level in fruit skin was ~2.14-, ~3.41-, ~5.25- and ~4.35-fold higher in G1, G2, G3 and G4, respectively. Briefly, anthocyanin content was lowest in G0 and highest in G3, followed by the contents in G4, G2, and G1. It was concluded that the more intense red colouring pattern in the apple pericarp of bud sport mutants was mainly caused by the accumulation of anthocyanins. In addition, a more blushed colouring pattern was observed with an increase in the number of bud sport generations. However, the third-generation mutant G3 showed greater blushing than did the fourth-generation mutant G4 at S1 and S2, and there was no significant difference at S3.

### The contents of IAA and ABA in apple pericarp increased with bud sport generation at veraison, while the content of GA decreased

Hormone levels were also analysed at three time points (S1, S2, and S3). The overall trend of IAA concentrations in apple pericarp first increased and then decreased from S1 to S3 and peaked at S2 (see Figure 1C). However, the GA content decreased from S1 to S3 (see Figure 1D). ABA concentrations of G0, G1 and G2 peaked at S1, whereas those of G3 and G4 peaked at S2 (see Figure 1E). Importantly, the IAA and ABA contents of G3 and G4 in apple pericarp at veraison S2 were considerably higher than those of G2, while G0 and G1 showed considerably lower levels than did G2. Nevertheless, GA concentrations, which displayed a trend opposite that of IAA and ABA, decreased with bud sport generation, that is, from G0 to G4.

### Transcriptomic profiling of the pericarp of ‘Red Delicious’ and its four continuous generation mutants

Triplicate sampling of the pericarp of ‘Red Delicious’ and its four continuous generation mutants at S2 yielded 15 RNA samples for RNA sequencing (RNA-seq) analysis, and the mapping rate of 20,189,457-26,765,510 clean reads onto the apple reference genome (https://iris.angers.inra.fr/gddh13/) (Daccord *et al*. 2017) ranged from 83.29% to 84.81% (Table 1). The average number of mapped reads ranged from 34,093,629 in G2 to 43,153,440.33 in G3. An FDR < 0.01 and fold changes ≥ 2 were used as screening criteria for DEGs; in addition, an FPKM (fragments per kilobase of mapped reads) ≥ 1.0 in at least one of the 5 triplicate samples was considered to be expressed. The mean normalized expression value (FPKM) per transcript of the three biological replicates was calculated for each sample using the geometric normalization method. The resulting dataset comprising 33,192 transcripts was used for subsequent analysis (see Dataset S1).

### Differentially expressed genes (DEGs) in ‘Red Delicious’ versus its mutants gradually increased with bud sport generation at veraison

To identify DEGs in ‘Red Delicious’ and its four generation mutants, seven pairwise transcriptome comparisons (i.e., G0 versus G1, G0 versus G2, G0 versus G3, G0 versus G4, G1 versus G2, G2 versus G3, and G3 versus G4) were performed at S2 (see Figure 2, Figure S1 and Table S1). The total number of DEGs was 3,466, including 1,456 up-regulated DEGs and 2029 down-regulated DEGs (see Figure 2 and Dataset S2). Among them, the number of both up-regulated and down-regulated DEGs increased considerably from the first generation mutant G1 to the fourth generation mutant G4 versus the number observed in G0, and the smallest number was observed in G1 versus that in G2. Moreover, we found that more genes were down-regulated than up-regulated in G1, G2, G3 and G4 relative to G0.

**Figure 2.**
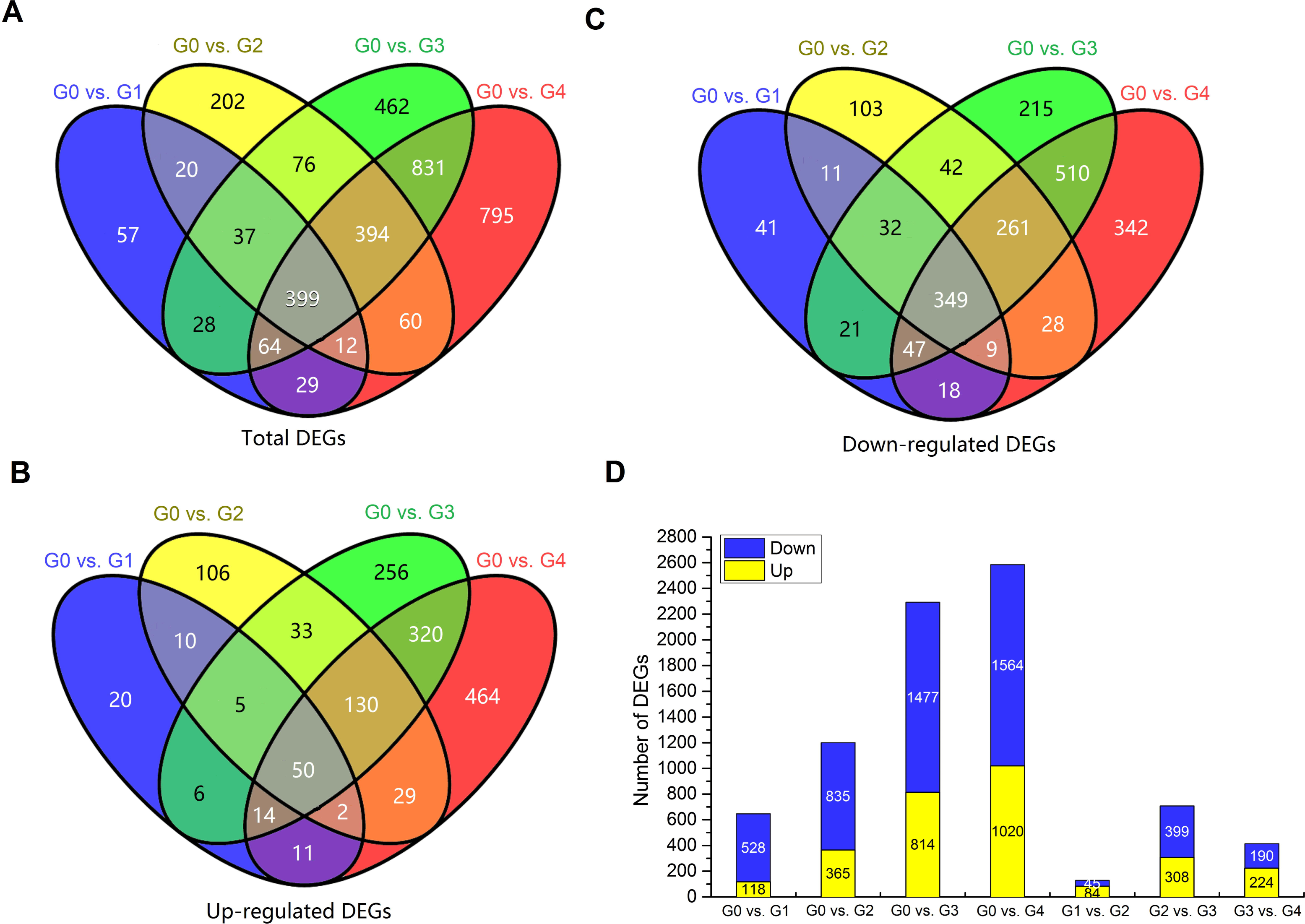
Summary of the number of differentially expressed genes (DEGs) identified by RNA-seq analysis in the pericarp tissues of ‘Red Delicious’ and its four generation mutants (‘Starking Red’, ‘Starkrimson’, ‘Campbell Redchief’ and ‘Vallee spur’) at S2, named G0 to G4. The number of total DEGs (A), up-regulated DEGs (B), and down-regulated DEGs (D) are presented by Venn diagrams (FDR < 0.01 and fold change ≥ 2). D, Number of total up-regulated and down-regulated DEGs. The histogram represents the number of commonly down-regulated (blue) and up-regulated (yellow) DEGs.

### Comparative transcriptome enrichment analysis identified key processes responsible for anthocyanin accumulation in ‘Red Delicious’ and its mutants

To understand the major functional categories represented by the DEGs, gene ontology (GO) enrichment analysis was carried out using all reference genes as background via the GOseq R package. GO term enrichment analysis categorized the annotated sequences into three main categories: biological process, cellular component and molecular function (see Table S2). In the biological process category, three significantly enriched terms, namely, “defence response to fungus”, “1-aminocyclopropane-1-carboxylate biosynthetic process” and “negative regulation of growth”, were shared in G0 versus G1, G0 versus G2, G0 versus G3 and G0 versus G4; among these terms, “1-aminocyclopropane-1-carboxylate biosynthetic process” and “negative regulation of growth” were also enriched in G1 versus G2, G2 versus G3 and G3 versus G4. Furthermore, in addition these two GO terms, three other significantly enriched GO terms, namely, “chitin catabolic process”, “regulation of leaf development” and “DNA conformation change”, were shared in G0 versus G1, G1 versus G2, G2 versus G3 and G3 versus G4. The cellular component category was further classified into “elongator holoenzyme complex”, which was enriched in G0 versus G1, G0 versus G2, G0 versus G3 and G0 versus G4, whereas, “elongator holoenzyme complex”, “mitochondrial intermembrane space” and “U12-type spliceosomal complex” were shared in G0 versus G1, G1 versus G2, G2 versus G3 and G3 versus G4. In the molecular function category, the DEGs were further classified into nine terms in the seven abovementioned comparison groups: “ADP binding”, “L-iditol 2-dehydrogenase activity”, “acyl-CoA hydrolase activity”, “3-beta-hydroxy-delta5-steroid dehydrogenase activity”, “O-methyltransferase activity”, “1-aminocyclopropane-1-carboxylate synthase activity”, “naringenin-chalcone synthase activity”, “catechol oxidase activity” and “trans-cinnamate 4-monooxygenase activity”. Moreover, “chitinase activity”, “sulfur compound binding” and “caffeate O-methyltransferase activity” were only observed in G0 versus G1, G1 versus G2, G2 versus G3 and G3 versus G4.

To further systematically understand the molecular interactions among the DEGs, we performed KEGG analysis using KOBAS software. The significantly enriched KEGG pathway term “sesquiterpenoid and triterpenoid biosynthesis” was shared in G0 versus G1, G0 versus G2 and G0 versus G3, but not in G0 versus G4 (Table 2). However, the “sesquiterpenoid and triterpenoid biosynthesis” pathway was derived from “terpenoid backbone biosynthesis”, which occurred both in G0 versus G3 and G0 versus G4. Furthermore, the number of DEGs belonging to the triterpenoid biosynthesis pathway gradually increased from G0 versus G1 to G0 versus G4. In addition, the “flavonoid biosynthesis” pathway was enriched in G0 versus G2, G0 versus G3 and G0 versus G4. Moreover, the “plant hormone signal transduction” pathway was enriched in G0 versus G1, G0 versus G2, G0 versus G3, G0 versus G4, G2 versus G3 and G3 versus G4 but not in G1 versus G2. Thus, the four abovementioned candidate pathways were considered to be heavily involved in anthocyanin accumulation.

### Functional classification of DEGs in ‘Red Delicious’ and its four continuous generation mutants

To further identify the major functions of DEGs and establish the pericarp pigment transcriptome, clustering analysis was applied to the 3,466 DEGs. These genes were grouped into six expression patterns (see Figure 3 and Dataset S3). Cluster 1 contained 561 DEGs whose expression peaked at G0/G1/G2. Cluster 2 contained 336 DEGs whose expression peaked at G3/G4. Cluster 3 contained 363 DEGs whose expression peaked at G2. Furthermore, 1049 and 348 DEGs whose expression peaked at G0 were included in clusters 4 and 5, respectively. Cluster 6 contained 809 DEGs whose expression peaked at G4. KEGG analysis was also carried out for DEGs belonging to each pattern with a *P*-value ≤ 0.01. The expression pattern of cluster 2 was positively consistent with total anthocyanin content (see Figure 1B), whereas clusters 4 and 5 were negatively aligned. DEGs in cluster 2 were significantly enriched in “plant hormone signal transduction”, “flavonoid biosynthesis”, “flavone and flavonol biosynthesis”, “phenylalanine metabolism”, and “phenylpropanoid biosynthesis”. Among those, some of the final metabolites of “flavonoid biosynthesis”, “flavone and flavonol biosynthesis”, “phenylalanine metabolism”, and “phenylpropanoid biosynthesis” are anthocyanins (Massonnet *et al*. 2017). Remarkably, the pathway that was co-enriched by clusters 4 and 5 was “sesquiterpenoid and triterpenoid biosynthesis”, which was hypothesized to be negatively related to the accumulation of anthocyanins. In addition, the pathway of “terpenoid backbone biosynthesis” was also enriched in cluster 1, and “flavonoid biosynthesis” and “phenylalanine metabolism” were enriched in cluster 3. Overall, the results of the functional classification analysis of the common expression patterns of DEGs combined with KEGG agreed with the results of the aforementioned comparative transcriptome enrichment analysis. Therefore, pathways including “flavonoid/phenylpropanoid biosynthesis”, “terpenoid biosynthesis” and “plant hormone signal transduction” were selected for subsequent analysis.

**Figure 3.**
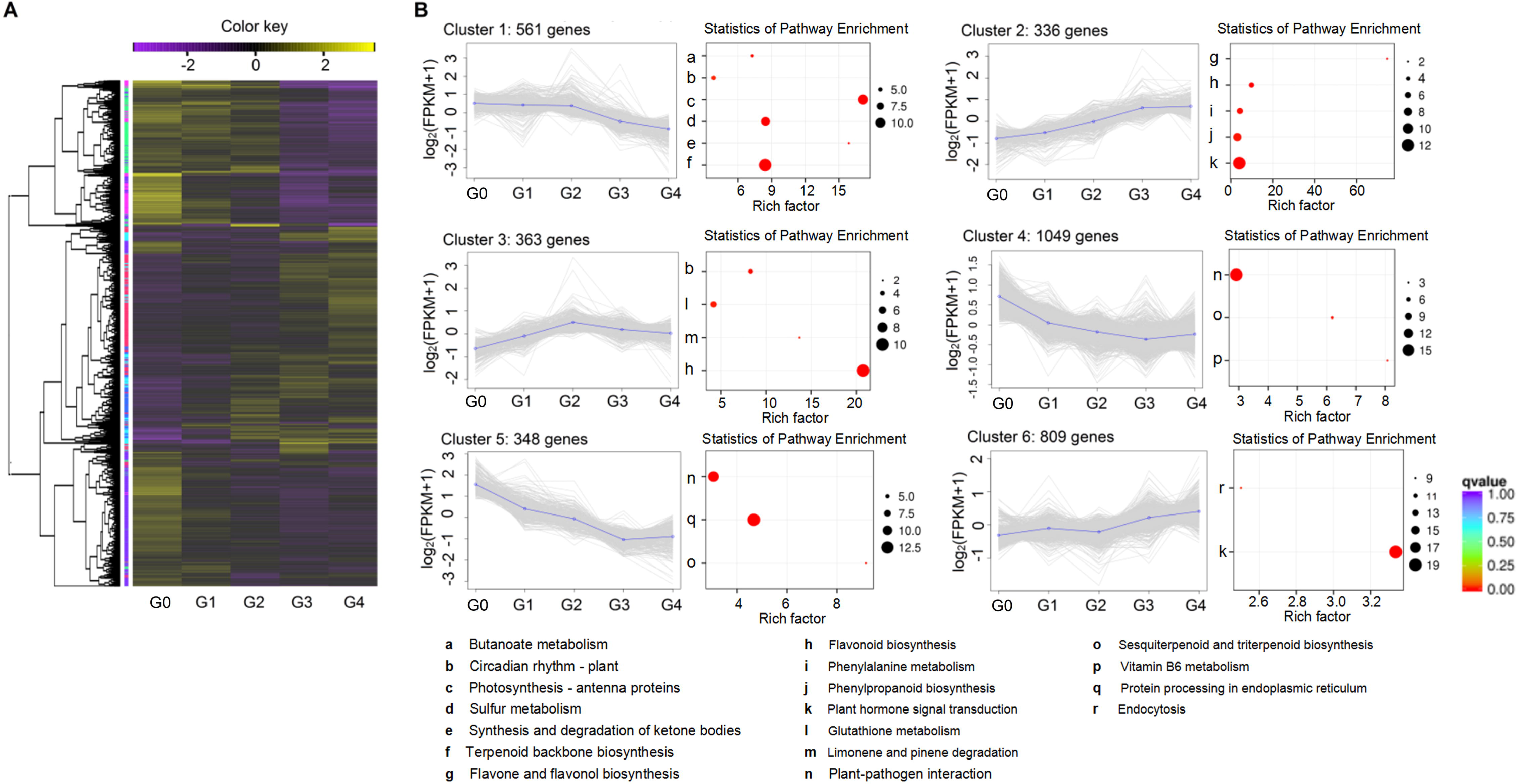
Gene expression profiles and KEGG category distribution of the DEGs in the six common expression clusters composing the pericarp transcriptome at S2. Clusters were derived by coupled clustering analysis of the 3,466 commonly modulated genes. A, Heat map of the overall common expression pattern. B, Each line represents the log_2_-transformed average of the mean FPKM values for an individual transcript. Significantly overrepresented KEGG categories are represented by red dots. KEGG category enrichment was computed using the R language, Cluster package, Biobase package, and *Q*-value package (*P* ≤ 0.01).

### Key candidate DEGs responsible for anthocyanin accumulation in ‘Red Delicious’ and its mutants

#### Genes involved in phenylpropanoid/flavonoid biosynthesis pathway

Variety-specific trends in the expression of phenylpropanoid/flavonoid biosynthesis pathway genes at S2 were investigated by preparing heat maps (see Figure 4A). We focused on the 28 DEGs involved in this pathway, including 4 from cluster 1, 13 from cluster 2, 10 from cluster 3, and 1 from cluster 5 (Table 3). Differences in the expression pattern of these genes from cluster 2 were found among ‘Red Delicious’ and its four continuous generation mutants, closely mirroring the differences in total anthocyanin concentration at S2. For example, all DEGs involved in the phenylpropanoid biosynthesis pathway, including one aspartate aminotransferase cytoplasmic (*ASP3*), one Phe ammonia lyase (*PAL*), two beta-glucosidase (*BGLU*), one 4-coumarate-CoA ligase (*4CL*) and three peroxidases (*PER*), were from cluster 2, demonstrating a gradually increasing expression pattern from G0 to G4 (see Figure 4A and Dataset S4). Among the DEGs, *ASP3* participates in the synthesis of phenylalanine and phenylpyruvate, which are precursors of anthocyanin synthesis. *BGLU* and *PER* are involved in coumarine and lignin biosynthesis, respectively. *PAL* and *4CL* are phenylpropanoid genes. In addition, down-regulated genes, including two quinate hydroxycinnamoyl transferase (*HCT*) and one caffeoyl-CoA O-methyltransferase (*CCoAOMT*), and up-regulated genes, including three cytochrome P450 98A2-like (*CYP98A*), four *CHS*, one avanone-3-hydroxylase (*F3H*), one dihydroflavonol reductase (*DFR*), and one anthocyanidin synthase (*ANS*), are involved in the synthesis of delphinidin from the phenylpropanoid biosynthesis pathway. Furthermore, the four *CHS* genes described above, one down-regulated and one up-regulated *CHI* gene, two *F3’H* genes, and the aforementioned *F3H*, *DFR* and *ANS* are involved in the synthesis of cyanidin. *F3’H* and other genes involved in pelargonidin synthesis are the same as those involved in cyanidin synthesis. These findings confirm that delphinidin, cyanidin and pelargonidin are the three main substances responsible for the synthesis of anthocyanins via the phenylpropanoid biosynthesis pathway in apple. Moreover, two down-regulated *FLS* genes are involved in flavone and flavonol biosynthesis.

**Figure 4.**
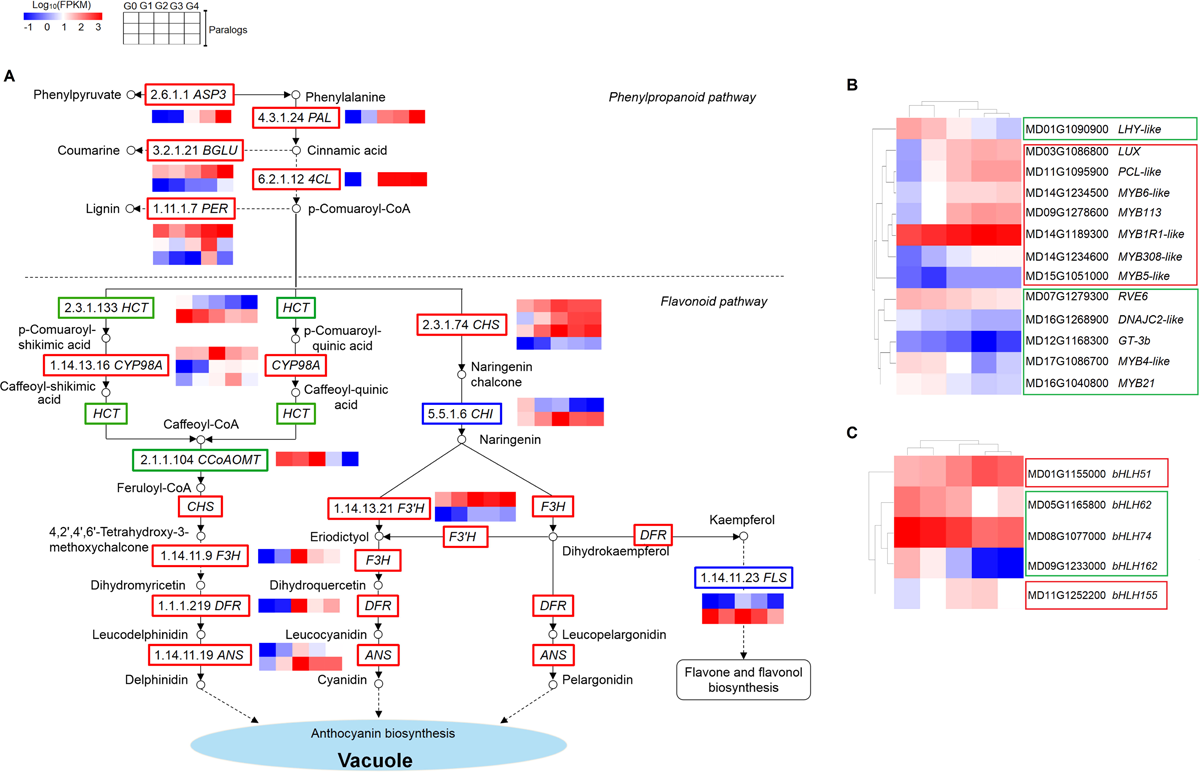
A, Differential expression of genes involved in phenylpropanoid/flavonoid biosynthesis pathway hormone signal transduction pathway in ‘Red Delicious’ and its four generation mutants (‘Starking Red’, ‘Starkrimson’, ‘Campbell Redchief’ and ‘Vallee spur’), named G0 to G4. Differential expression of transcription factors that encode myb-like (B) and helix-loop-helix (C) DNA-binding domains. Heat maps depict the normalized gene expression values, which represent the means ± SD of three biological replicates. Expression values of five libraries are presented as FPKM normalized log_10_-transformed counts.

#### Genes involved in myb-like and helix-loop-helix DNA-binding domain transcriptional factors

Correlative analysis was carried out between anthocyanin content at S2 and the expression levels of transcription factors that encode myb-like and helix-loop-helix DNA-binding domains in DEGs (see Dataset S5, 6). As a result, 13 MYB and 5 bHLH transcription factors were screened. Among them, the expression of MYB members, including *LUX*, *MYB113*, *PCL*-*like*, *MYB1R1*-*like*, *MYB6*-*like*, *MYB308*-*like* and *MYB5*-*like*, bHLH members, including *bHLH51* and *bHLH155*, presented a notable positive correlation with anthocyanin content at S2 (*P*<0.05) (see Figure 4B, C). MYB members *LHY-like*, *RVE6*, *GT-3b*, *MYB21*, DNAJC2-like and *MYB4-like* and bHLH members *bHLH62*, *bHLH74* and *bHLH162* showed a remarkably negative correlation with anthocyanin content at veraison S2 (*P*<0.05).

#### Genes involved in terpenoid biosynthesis and plant hormone signal transduction

KEGG analysis showed that terpenoid backbone biosynthesis was over-presented in cluster 1 and that sesquiterpenoid and triterpenoid biosynthesis branching from terpenoid backbone biosynthesis was over-presented in clusters 4 and 5 (see Figure 3), indicating that this pathway may play important roles in anthocyanin accumulation in apple. Moreover, the plant hormone signal transduction pathway, which is modulated by terpenoid biosynthesis, was enriched in cluster 2. A total of 18 DEGs in the terpenoid biosynthesis pathway (see Dataset S7 and Table 4) and 12 DEGs in the plant hormone signal transduction pathway (see Dataset S8 and Table 5) were also investigated by preparing heat maps (see Figure 5). The expression of DEGs involved in terpenoid backbone biosynthesis, including one acetyl-CoA acetyltransferase, cytosolic 1 (*AACT1*), two hydroxymethylglutaryl-CoA synthase-like (*HMGS*), two 3-hydroxy-3-methylglutaryl-coenzyme A reductase 1-like (*HMGR*), one mevalonate kinase-like (*MVK*), two diphosphomevalonate decarboxylase MVD2-like (*MVD2*), one isopentenyl-diphosphate Delta-isomerase I (*IDI1*), three farnesyl pyrophosphate synthase 2-like (*FPPS2*) and six squalene monooxygenase-like (*SQMO*), were gradually down-regulated from G0 to G4. Nevertheless, two auxin transporter-like 1 (*AUX1*), four auxin-responsive protein *AUX/IAA* and three auxin-responsive protein *SAUR* involved in tryptophan metabolism of auxin biosynthesis; one F-box protein GID2-like (*GID2*) involved in diterpenoid biosynthesis of GA biosynthesis; and one protein phosphatase 2C (*PP2C*) and one sucrose non-fermenting-1-related protein kinase 2 (*SnRK2*) involved in carotenoid biosynthesis of ABA biosynthesis were up-regulated from G0 to G4.

To further evaluate the validity of our results, 16 representative DEGs used previously as shown in Figures 4 and 5 were selected for expression level examination by qRT-PCR (see Table S3). The overall trend of relative expression levels at three stages was consistent with that of deep sequencing at S2 (see Figure 6), suggesting that the candidate genes involved in pathways such as phenylpropanoid/flavonoid biosynthesis, terpenoid biosynthesis and plant hormone signal transduction, appended with MYB and bHLH transcriptional factors, were directly correlated with anthocyanin accumulation.

**Figure 5.**
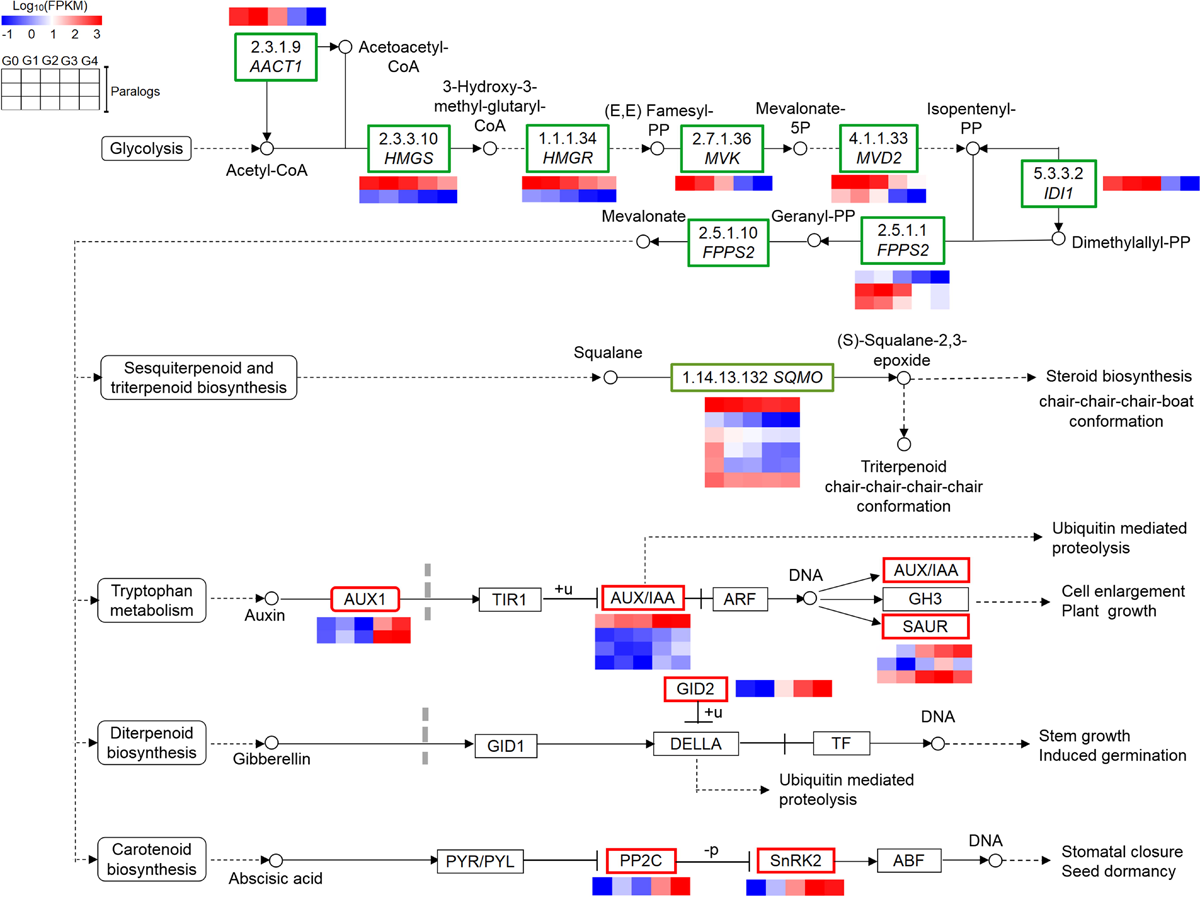
Differential expression of genes involved in sesquiterpenoid and triterpenoid/terpenoid biosynthesis coupled with hormone signal transduction pathway in ‘Red Delicious’ and its four generation mutants (‘Starking Red’, ‘Starkrimson’, ‘Campbell Redchief’ and ‘Vallee spur’), named G0 to G4. Heat maps depict the normalized gene expression values, which represent the means ± SD of three biological replicates. Expression values of five libraries are presented as FPKM normalized log_10_-transformed counts.

**Figure 6.**
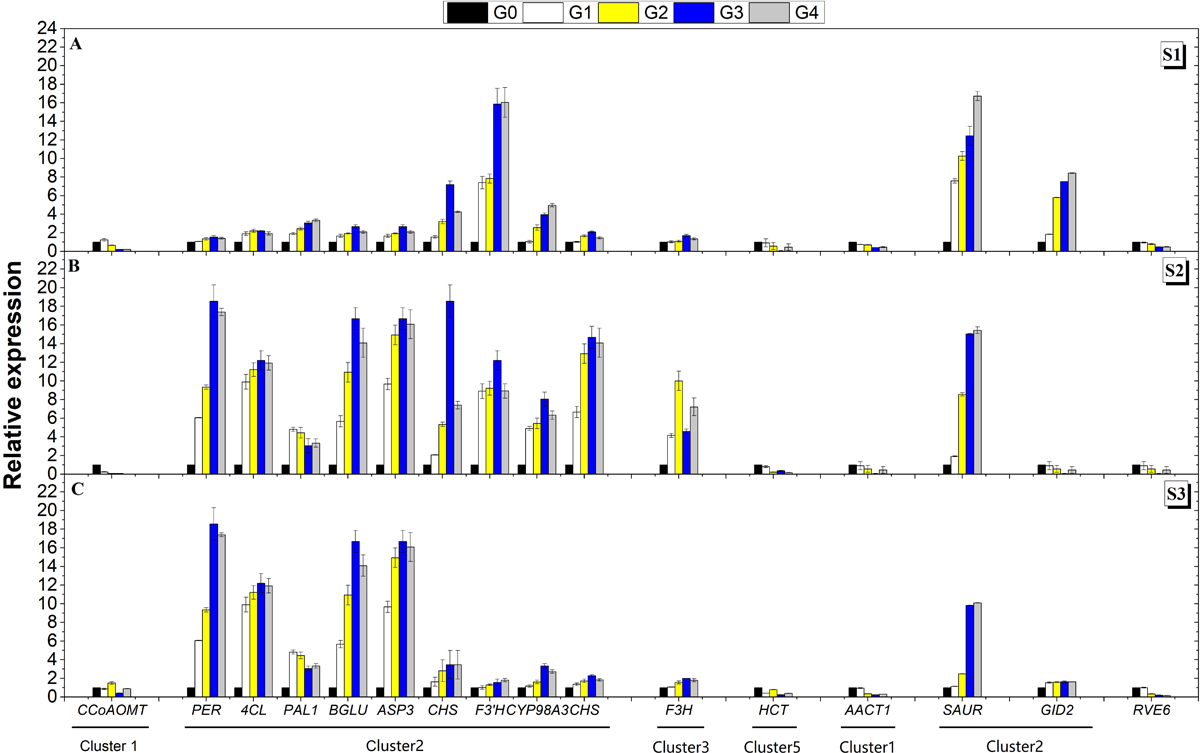
Relationships between total anthocyanin contents and transcript levels of the sixteen representative genes from Figures 4 and 5. For each accession, the expression was determined in three developmental stages (S1 to S3) of pericarp tissues. For the qRT-PCR assay, the mean was calculated from three biological replicates, each with three technical replicates (*n*=9). These replicates were then normalized relative to the expression of *MdGADPH*. The x-axis in each chart is the same and represents different Malus accessions, as indicated by names in the bottom panel, which are arranged in different clusters. The left y-axis represents relative expression levels determined by qRT-PCR.

## DISCUSSION

### The activation of early phenylpropanoid biosynthesis genes was more responsible for anthocyanin accumulation in apple pericarp of bud sport mutants

In plants, phenylpropanoid biosynthesis gives rise to a large number of secondary metabolites, including hydroxycinnamic acids, monolignols/lignin, coumarins, benzoic acids, stilbenes, anthocyanins and flavonoids, serving different functions in plant development, reproduction, defence, and protection against biotic/abiotic stresses (Voge 2010; Zhang *et al*. 2013). Differences in the expression pattern of genes involved in phenylpropanoid/flavonoid biosynthesis result in diverse anthocyanin profiles (El-Sharkawy *et al*. 2015; Massonnet *et al*. 2017). Our survey provided a comprehensive profile of the phenylpropanoid/flavonoid biosynthesis pathway in ‘Red Delicious’ and its four continuous generation mutants. The results showed that all of the early phenylpropanoid biosynthesis pathway genes, including *ASP3*, *PAL*, *4CL*, *BGLU* and *PER*, were aggregated in cluster 2 (see Figure 4A and Table 3), which matched the anthocyanin content (see Figure 1A, B). Other genes in cluster 2 containing *CHS*, *CYP98A* and *F3’H* are involved in the middle steps of the phenylpropanoid pathway, that is, the early steps of the flavonoid biosynthesis pathway. Nevertheless, genes encoding *HCT*, *CCoAOMT*, *CHS*, *CYP98A*, *CHI*, *F3H*, *DFR*, *FLS* and *ANS* were involved in the middle and late steps of the phenylpropanoid biosynthesis pathway and gathered in clusters 1 and 3. Thus, the activation of early phenylpropanoid biosynthesis pathway genes was demonstrated to be most responsible for pigment accumulation in the apple pericarp of bud sport mutants. In addition, *ASP3*, *BGLU* and *PER* were confirmed to be involved in the synthesis of phenylpyruvate, coumarine and lignin, respectively (see Figure 4A). Interestingly, 44 stilbene synthase (*STS*) genes involved in stilbene biosynthesis were characterized to influence anthocyanin accumulation during grapevine (*Vitis vinifera*) maturation by Massonnet *et al*. (2017). Nevertheless, these genes do not exist among our 3,466 DEGs, possibly because of their variety-specific nature (Zenoni *et al*. 2016).

### MYB and bHLH modulated anthocyanin accumulation in apple pericarp by regulating the transcription of genes involved in the phenylpropanoid/flavonoid pathway

MYB and bHLH autonomously mediated the transcription of genes involved in the middle steps of the phenylpropanoid pathway, that is, the early steps of the flavonoid biosynthetic pathway (*CHS*, *CHI*, *F3H*, *F3’H* and *FLS*), which leads to the production of colourless dihydroflavonol compounds (Baudry *et al*. 2004; Stracke *et al*. 2007; Petroni and Tonelli 2011; Hu *et al*. 2016; An *et al*. 2017). The heterologous expression of OjMYB1 in Arabidopsis could enhance anthocyanin content and up-regulate the expression levels of structural gene-related anthocyanin biosynthesis (Feng *et al*. 2017). The red radish (*Raphanus sativus* L.) bHLH transcription factor RsTT8 acts as a positive regulator of anthocyanin biosynthesis (Lim *et al*. 2017). Nevertheless, *AtMYB4*, *AmMYB308*, *FaMYB1*, *ZmMYB31*, *ZmMYB42*, *PhMYB4, VvMYBC2-L1* and *PtrMYB57* have been demonstrated to repress phenylpropanoid synthesis, likely via repression of synthesis genes (Tamagnone *et al*. 1998; Aharoni *et al*. 2001; Colquhoun *et al*. 2011; Huang *et al*. 2014; Wan *et al*. 2017). We corroborated that MYB members, including *LUX*, *MYB113*, *PCL-like*, *MYB1R1-like*, *MYB6-like*, *MYB308-like* and *MYB5-like*, and bHLH members, including *bHLH51* and *bHLH155*, showed a notable positive correlation with anthocyanin content (see Dataset S5, 6 and Figure 4B, C) and were considered to promote anthocyanin synthesis by mediating the transcription of structural genes, *CHS* and *F3’H*, which are involved in the flavonoid pathway. Other MYB members, including *LHY-like*, *RVE6*, *GT-3b*, *MYB21*, *DNAJC2-like* and *MYB4-like*, and bHLH members, including *bHLH62*, *bHLH74* and *bHLH162* showed a remarkably negative correlation with anthocyanin content and were demonstrated to repress anthocyanin synthesis. In addition, HD-Zip I transcription factor MdHB1 was involved in the regulation of anthocyanin accumulation (Lü *et al*. 2014). When MdHB1 is silenced, *MdMYB10*, *MdbHLH3*, and *MdTTG1* are released to activate the expression of *MdDFR* and *MdUFGT* and anthocyanin biosynthesis, resulting in red flesh in apple cv. ‘Granny Smith’ (Jiang *et al*. 2017). The expression of *F3’5’H*, *DFR* and *ANS* is strongly inhibited by the increase in the expression of *MYBL1*, which is a novel R3 MYB transcription factor classified as an MYB transcriptional repressor (Gates *et al*. 2018). However, a full understanding of the mechanism by which structural genes involved in anthocyanin synthesis are specifically mediated by MYB and bHLH remains elusive clear and requires further investigation.

### Terpenoid biosynthesis modulated anthocyanin accumulation by positively regulating plant hormone signal transduction in apple pericarp of bud sport mutants

Hormones are important factors inducing anthocyanin accumulation (Jiang and Fu 2007; Shan 2009; Loreti *et al*. 2010). Carvalho *et al*. (2010) provided evidence that anthocyanin accumulation was promoted by exogenous ABA and CTK and inhibited by GA in tomato hypocotyls. Co-treatment of IAA and CTK significantly enhanced the cytokinin-induced increase in anthocyanin levels, but an auxin concentration that was too high strongly inhibited anthocyanin synthesis even in the presence of cytokinin in callus cultures of red-fleshed apple (*M. sieversii* f.*niedzwetzkyana*), as shown by Ji *et al*. (2014). Our results showed that sesquiterpenoid and triterpenoid biosynthesis along with plant hormone transduction, including tryptophan metabolism for IAA, diterpenoid biosynthesis for GA and carotenoid biosynthesis for ABA, branches from the general terpenoid backbone synthesis pathway and shares the same precursors as glycolysis (see Figure 5). The sesquiterpenoid and triterpenoid biosynthesis pathways comprised six *SQMO* genes in clusters 4 and 5, which were negatively correlated with anthocyanin content. The genes *AACT1*, *HMGS*, *HMGR*, *MVK*, *MVD2*, *IDI1* and *FPPS2*, involved in the early steps of terpenoid backbone synthesis, were in cluster 1, which also reflected the differential accumulation of anthocyanin to some extent. Moreover, *AUX1*, *AUX/IAA* and *SAUR*, associated with tryptophan metabolism for IAA; *GID2*, associated with diterpenoid biosynthesis for GA; and *PP2C* and *SnRK2*, associated with carotenoid biosynthesis for ABA, were in cluster 2, which positively reflected the anthocyanin content. Likewise, the contents of IAA and ABA increased, while GA decreased with maturation and pigment accumulation from G0 to the fourth-generation mutant G4 (see Figure 1C, E). Therefore, the down-regulation of genes involved in terpenoid biosynthesis positively induced the expression of *AUX1*, *AUX/IAA*, *SAUR*, *GID2*, *PP2C* and *SnRK2*, resulting in increased synthesis of IAA and ABA and decreased synthesis of GA, thus modulating anthocyanin accumulation. Loreti *et al*. (2010) suggested the existence of crosstalk between the sucrose and hormone signalling pathways in the regulation of the anthocyanin biosynthetic pathway. Similarly, exogenous ABA promoted fruit ripening by increasing anthocyanin content in sweet cherry (*Prunus avium* L.) cv. Sato Nishiki, and the expression of *PaPP2C3*, *PaPP2C5* and *PaPP2C6* was significantly induced by exogenous ABA (Wang *et al*. 2015). In general, there may be some form of crosstalk between the activation of phenylpropanoid biosynthesis and plant hormone signal transduction in pigment accumulation of apple bud sport mutants.

## CONCLUSIONS

We investigated the pericarp transcriptome of ‘Red Delicious’ and its four continuous generation mutants (‘Starking Red’, ‘Starkrimson’, ‘Campbell Redchief’ and ‘Vallee spur’) and identified specific processes that lead to the accumulation of anthocyanin. The results indicate that apple pericarp pigmentation and anthocyanin content were increased in the mutants due to bud sport. Terpenoid biosynthesis influences anthocyanin accumulation by positively regulating the synthesis of IAA and ABA and negatively regulating the synthesis of GA. MYB and bHLH modulate anthocyanin accumulation in apple pericarp by regulating the transcription of genes *CHS* and *F3’H*. *ASP3*, *CYP98A* and *CCoAOMT* are novel anthocyanin-associated genes of apple first reported the present study. This novel set of genes provides not only new insights into anthocyanin biosynthesis but also important clues for more dedicated studies to broaden our knowledge of the anthocyanin pathway in apple.

## ACKNOWLEDGMENTS

This research was financially supported by the Key Scientific Technology Research Projects of Gansu Province (GPCK2013-2), the Fostering Foundation for the Excellent Ph.D. Dissertation of Gansu Agricultural University (2017002), and the Science and Technology Major Project of Gansu Province (18ZD2NA006).

## Author contributions

Bai-Hong Chen designed the research. Wen-Fang Li carried out the experiments with the help of Juan Mao, Shi-Jin Yang, Zhi-Gang Guo, Zong-Huan Ma, Cun-Wu Zuo and Ming-Yu Chu. Wen-Fang Li collected the experimental data and drafted the manuscript. Mohammed Mujitaba Dawuda reviewed the manuscript and part of the data analysis. All authors read and approved the final manuscript.

## Supporting information

Additional Supporting Information may be found online in the supporting information tab for this article:

**Figure S1** Summary of the number of differentially expressed genes (DEGs) identified by RNA-seq analysis in the pericarp tissues of ‘Red Delicious’ and its four generation mutants (‘Starking Red’, ‘Starkrimson’, ‘Campbell Redchief’ and ‘Vallee spur’) at S2, named G0 to G4. The number of total DEGs (A), up-regulated DEGs (B), and down-regulated DEGs (D) are presented by Venn diagrams (FDR < 0.01 and fold change ≥ 2). D, Number of total up-regulated and down-regulated DEGs. The histogram represents the number of commonly down-regulated (blue) and up-regulated (yellow) DEGs.

**Table S1** Summary of RNA-Seq data and mapping metrics

**Table S2** Gene ontology (GO) enrichment analyses for DEGs in ‘Red Delicious’ and its four generation mutants.

**Table S3** Sequence of primers used for qRT-PCR analysis.

